# Increased expression of developmental Nav1.3 promotes hippocampal CA3 hyperexcitability in early-stage 5xFAD mice

**DOI:** 10.64898/2026.07.28.741298

**Authors:** Jiabin Tang, Cameron Swope, Kishan Patel, Azliana Jusnida Ahmad Jafri, Jiaxin Xiang, Teresa A. Milner, Hugh C. Hemmings, Jimcy Platholi

## Abstract

Neuronal hyperexcitability is an early and pervasive feature of Alzheimer’s disease (AD) that both predicts and accelerates subsequent cognitive decline. Persistent excitability depends on activation of voltage-gated sodium channels (Nav), yet most work has focused on the Nav subtypes expressed in the mature brain. Here, we show that Nav1.3, a subtype normally confined to early development, shows aberrantly increased expression in the dentate gyrus (DG)-CA3 circuit in early-stage 5xFAD mice. Using *in vivo* fiber photometry, three-month-old 5xFAD mice exhibited greater CA3 neuron population activity than seven-month-old 5xFAD mice or wildtype littermates. Oligomeric Aβ expression, assessed by immunolabeling, was sparse at three months and rose significantly by seven months, indicating that CA3 hyperactivity emerges before substantial oligomeric Aβ accumulates. This early activity increase coincided with elevated Nav1.3 expression at the DG-CA3 mossy fiber synapse, localized by immuno-electron microscopy to presynaptic mossy fiber terminals, where it exceeded levels in age-matched controls. Lentiviral shRNA-mediated knockdown of Nav1.3 expression in CA3 normalized CA3 network activity in early-stage 5xFAD mice. These findings identify Nav1.3 expression at DG-CA3 mossy fiber synapses as a driver of early hippocampal network dysfunction in AD and suggest Nav1.3 modulation as a potential target for circuit-level intervention.

## Introduction

Aberrant network activity is among the earliest detectable abnormalities in Alzheimer’s disease (AD), which emerges years before overt neurodegeneration. Functional imaging and electrophysiological recordings in individuals with mild cognitive symptoms reveal abnormal network synchronization (1, 2), and subclinical epileptiform activity is detected in a substantial proportion of patients by extended monitoring (3–5). This network dysfunction is increasingly recognized as a potentially causal feature of the disease rather than a secondary symptom (6). Defining the cellular and circuit mechanisms that drive this early hyperexcitability is essential both for understanding pathogenesis and for developing targeted therapies to slow progression and cognitive decline.

As AD progresses, vulnerability is region- and cell-type-specific, with distinct neuronal populations affected at different stages (7–9). The hippocampus is central to this process, and alterations across the dentate gyrus (DG)-CA3 pathway are particularly informative for early disease (10, 11). CA3 is uniquely predisposed to pathological hyperactivity: its recurrent excitatory architecture and intrinsic conductances support hypersynchronous outbursts and persistent, action potential-dependent firing (12–14). Several AD models report abnormal CA3 activity preceding overt pathology, implicating CA3 dysregulation as an early, potentially driving event (15–17). Notably, *in vivo* imaging indicates that soluble Aβ can be sufficient to trigger early hippocampal hyperactivity (18), although the circuit- and synapse-level substrates that render specific hippocampal nodes vulnerable remain incompletely defined. A principal determinant of CA3 excitability is the powerful excitatory drive supplied by DG granule cells through large, specialized mossy fiber terminals (MFTs) (19, 20). MFTs are multi-release-site boutons that directly innervate and selectively drive CA3 pyramidal neurons, so molecular or functional changes at this synapse can profoundly reshape network synchronization. As the mossy fiber synapse both governs physiological ensemble recruitment and is sensitive to developmental and injury-induced reprogramming (21), synapse-specific changes here are a plausible mechanism linking early molecular pathology to the aberrant CA3 network states.

Voltage-gated sodium channels (Nav) are principal regulators of neuronal excitability and critically shape network firing (22). Four major Nav subtypes, Nav1.1, Nav1.2, Nav1.3 and Nav1.6, are expressed in the mammalian brain (23, 24), each with characteristic cell-type and subcellular distributions. Nav1.1 is enriched in GABAergic interneurons, whereas Nav1.2 and Nav1.6 predominate in excitatory neurons and at axon initial segments (25–28). Loss or mutation of specific subtypes causes spontaneous seizures and severe childhood epilepsies (29, 30), and shifts in subtype composition can profoundly alter excitation-inhibition balance. In AD models, interneuron Nav1.1 loss and amyloid precursor protein (APP)-driven Nav1.6 upregulation have each been linked to network hyperexcitability (31–36) but these changes do not fully account for the early, cell-selective increases in excitatory drive seen in prodromal disease.

Nav1.3 is predominantly expressed during early brain development and is downregulated in the mature CNS (37, 38), although it can be re-expressed after injury and in other conditions associated with hyperexcitability (39–41). It is distinguished by a low activation threshold, rapid recovery from inactivation, and a capacity to sustain high-frequency firing (37, 42). These properties make Nav1.3 a strong candidate for driving early network changes in AD, particularly at mossy fiber synapses. Despite this pro-excitatory profile, whether Nav1.3 is pathologically increased in the AD brain, and whether it contributes to early network dysfunction, has not been examined.

Using the 5xFAD model of AD together with *in vivo* fiber photometry, electron-microscopic immunolocalization, and lentiviral-mediated knockdown, we show that Nav1.3 shows aberrantly increased expression at the DG-CA3 mossy fiber synapse in AD mice relative to age-matched controls, coincident with peak CA3 hyperactivity at three months when oligomeric Aβ remains low. Lentiviral-mediated knockdown of Nav1.3 in CA3 normalizes CA3 network activity in early AD. These findings identify developmental Nav1.3 expression at DG-CA3 synapses as a mechanism driving pathological network activity in early AD and point to subtype-selective Nav modulation as a potential therapeutic strategy.

## Results

### Early-stage 5xFAD mice show elevated hippocampal CA3 neuronal activity

Hippocampal hyperexcitability emerges early in AD and may contribute to cognitive decline (5, 43, 44). To test whether CA3 activity is elevated at early disease stages, we recorded calcium dynamics by fiber photometry from dorsal CA3 in male and female 3- and 7-month-old 5xFAD and wild-type (WT) mice expressing GCaMP6s (n= 13 per group; Fig. 1a-c). We pre-specified two comparisons: 3-month 5xFAD *vs* age-matched WT, and 3-month *vs* 7-month 5xFAD. Two-way ANOVA revealed a significant genotype x age interaction (Fig. 1c), alongside significant main effects of genotype and age; planned comparisons showed that CA3 activity was higher in 3-month 5xFAD than in both age-matched WT and 7-month 5xFAD, with no genotype difference at 7 months. The same directional pattern was present in both sexes but arose through different statistical signatures: significant genotype and age main effects in females, and a significant genotype x age interaction in males. In both sexes, planned comparisons confirmed elevated 3-month 5xFAD activity (Supplementary Fig. 1). Thus, CA3 hyperactivity is present early in 5xFAD mice of both sexes, with sex-specific differences in how genotype and age contribute to the effect.

**Figure 1.**
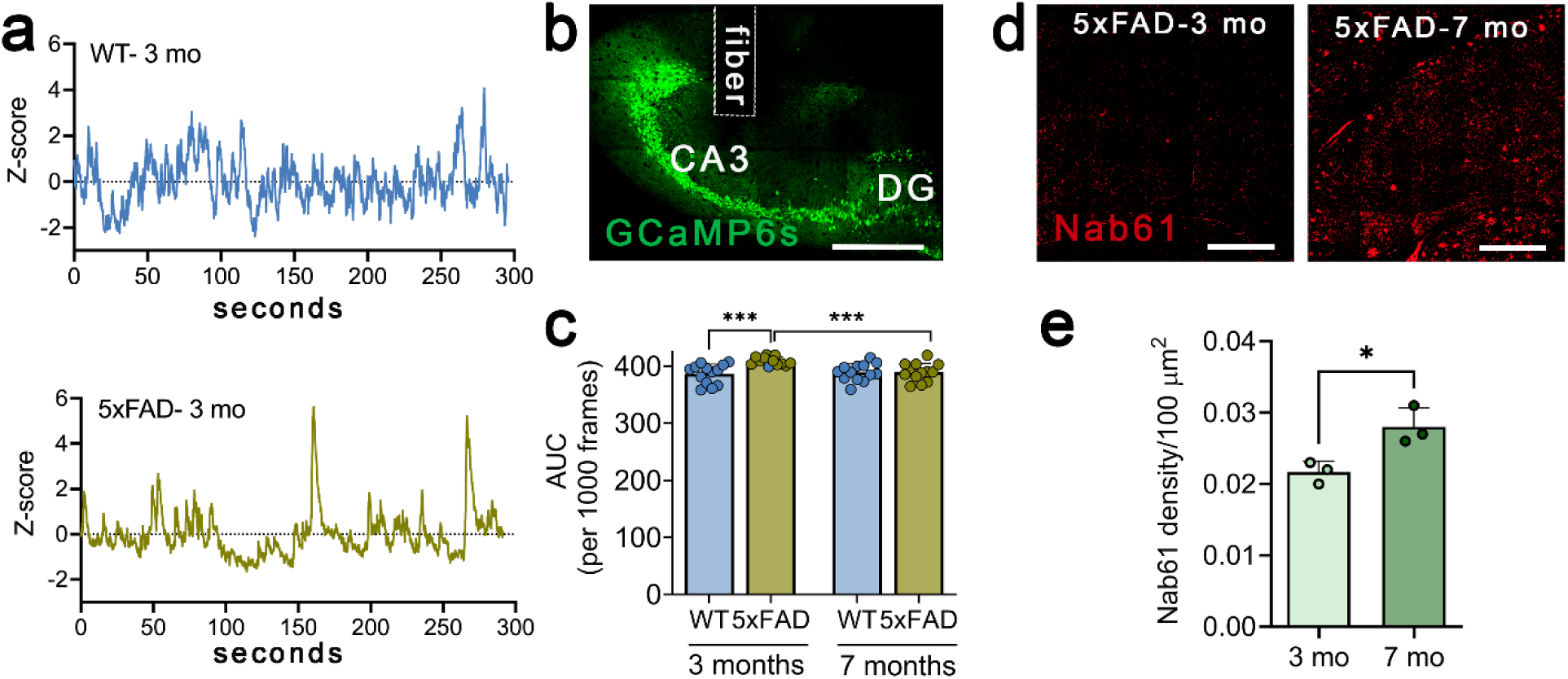
CA3 hyperactivity in early-stage 5xFAD mice. **a)** Example Z-scored (ΔF/F) fiber photometry traces from wildtype (top) and 5xFAD (bottom) mice at 3 months of age. **b)** Representative image of GCaMP6s expression and fiber placement in dorsal hippocampal CA3 region. Scale bar, 400 μm **c)** Quantification of mean CA3 activity (area under the curve (AUC) per 1000 frames) across groups. n = 13 mice per group. Data are mean ± SD. Full model two-way ANOVA (genotype x age) revealed a significant genotype x age interaction (F(1, 48) = 7.80, p= 0.0075), with significant main effects of age (F(1, 48) = 4.72, p = 0.035) and genotype (F(1, 48) = 8.44, p= 0.0055). Pre-specified planned comparisons (Sidak’s corrected) showed higher CA3 activity in 3-month 5xFAD than 3-month WT (p = 0.0002; mean difference 22.7 [95% CI 11.4-34.1] and than 7-month 5xFAD (p = 0.0010; mean difference 19.8 [95% CI 8.5-31.2] **d)** Representative images of Nab61^+^ Aβ oligomers in hippocampal CA3 from 3-month (left) and 7-month (right) 5xFAD mice. Scale bar, 400 μm **e**) Quantification of Nab61+ immunoreactivity (density/100 μm^2^) in 3-month and 7-month 5xFAD mice. Data are mean ± SD; individual points shown; n = 3 male mice per group. Two-tailed unpaired t-test; t (4) = 3.6, p = 0.023.

To test whether CA3 hyperactivity precedes oligomeric Aβ accumulation, we quantified Aβ oligomers by Nab61 immunofluorescence in male mice (Fig. 1d-e; n = 3 per group). Nab61-positive oligomers were scarce at 3 months but markedly increased by 7 months in 5xFAD hippocampus (Fig. 1d-e). Together with the photometry data, these results indicate that CA3 hyperexcitability is an early feature of the 5xFAD model that emerges prior to substantial Aβ oligomer deposition.

### Nav1.3 expression is increased in early-stage 5xFAD mice

The immature brain is relatively hyperexcitable in part because it expresses distinct voltage-gated sodium channel subtypes (38). Nav1.3 differs from adult subtypes by activating at a lower threshold and recovering from inactivation more rapidly, which support sustained, persistent firing (45–48). Although Nav1.3 is normally downregulated after development, it can be re-expressed after injury (49). To ask whether Nav1.3 expression is increased during neurodegeneration and whether this increase is associated with CA3 hyperactivity, we performed double-label immunofluorescence for Nav1.3 and the activity marker c-Fos in dorsal CA3 (Fig. 2a) and quantified both c-Fos^+^ cell density and Nav1.3/c-Fos colocalization. The density of c-Fos^+^ cells was significantly elevated in 3-month-old 5xFAD mice compared with age-matched WT controls and with 7-month-old 5xFAD mice (Fig. 2b). Analysis of Nav1.3/c-Fos colocalization showed a higher fraction of c-Fos^+^ neurons co-expressing Nav1.3 in 3-month-old 5xFAD mice relative to WT controls, whereas no genotype difference was observed at 7 months (Fig. 2c). The genotype x age interaction for colocalization was not significant, but there was a significant main effect of genotype. These findings suggest that increased Nav1.3 expression is enriched in activated CA3 neurons at an early disease stage in 5xFAD mice, consistent with a mechanistic link between increased Nav1.3 expression and hippocampal hyperexcitability.

**Figure 2.**
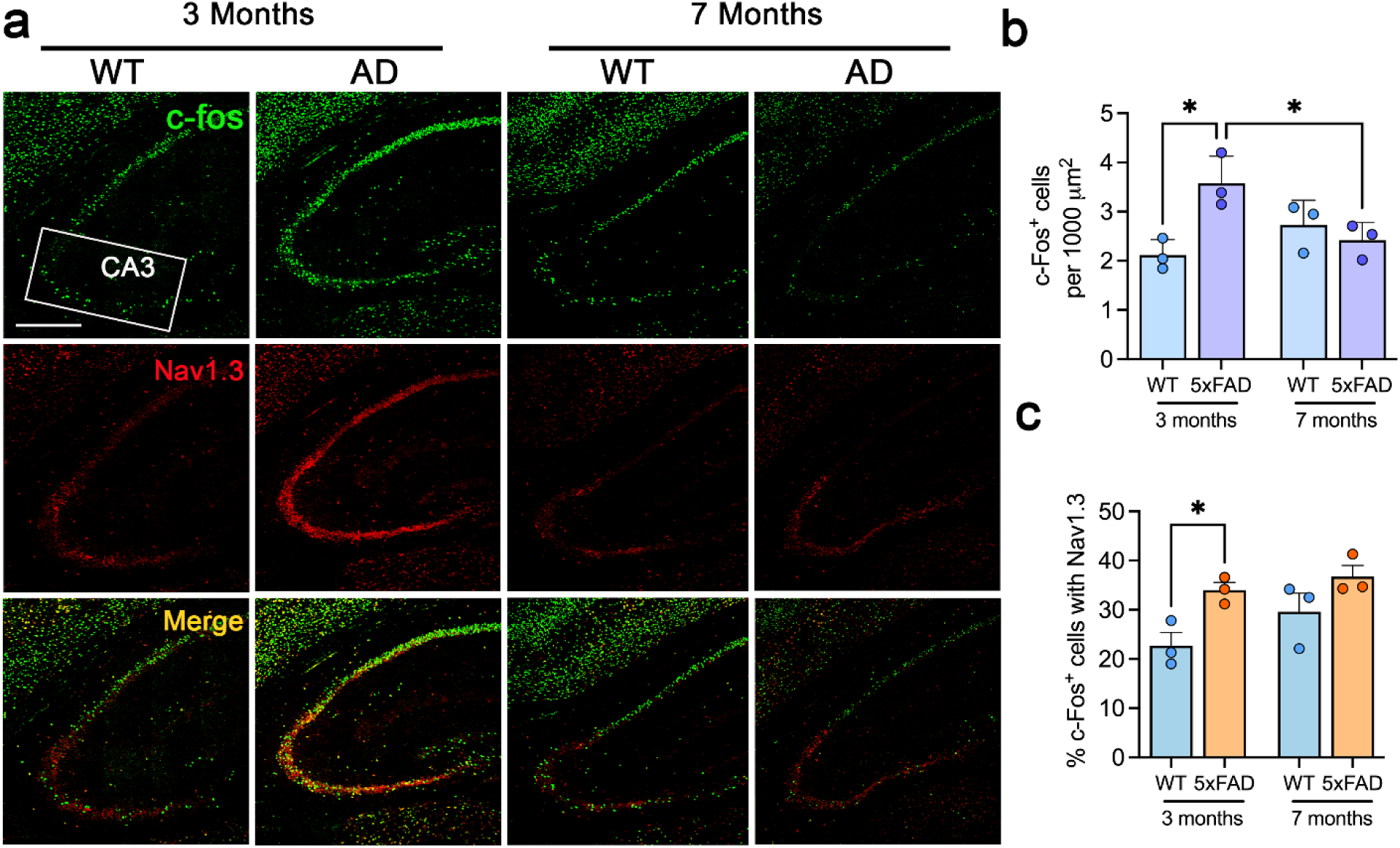
Nav1.3 expression is increased in activated CA3 neurons in early-stage 5xFAD mice. **a)** Representative confocal images of dorsal hippocampus CA3 showing c-Fos (green, top), Nav1.3^+^ (red, middle), and merged channels (yellow, bottom) for WT and 5xFAD at 3 months (left) and 7 months (right). The inset on the top-left panel indicates the CA3 subregion delineated using the Allen Brain Atlas where c-Fos/Nav1.3 were quantified. Scale bar = 400 μm. **b)** Quantification of neuronal activation in CA3, shown as c-Fos^+^ cells per 1000 μm^2^. c-Fos expression was higher in 3-month 5xFAD mice than in 3-month WT mice or 7-month 5xFAD mice (two-way ANOVA genotype x age interaction: F(1, 8) =12.0, p = 0.009; *post-hoc* Sidak: 5xFAD 3 months vs WT 3 months, p =0.015; 5xFAD 3 months vs 5xFAD 7 months, p = 0.049). **c)** Quantification of Nav1.3/c-Fos colocalization, shown as the percentage of c-Fos^+^ neurons that are Nav1.3^+.^ The genotype x age interaction was not significant for colocalization, but there was a significant main effect of genotype, with higher Nav1.3/c-Fos co-expression at 3 months in 5xFAD mice *vs* age-matched WT controls (F(1, 8) = 11.8, p = 0.009); *post-hoc* Sidak’s comparisons indicate increased colocalization in 3-month 5xFAD mice relative to 3-month WT (p = 0.035). No other comparisons reached significance. Data are mean ± SD; n = 3 mice per group, 9 fields analyzed per mouse.

Having established that Nav1.3 expression is increased in activated CA3 neurons early in AD pathology, we analyzed whether this increase occurs at synaptic sites capable of directly influencing CA3 excitability. MFTs from DG granule cells provide powerful excitatory input to CA3 and are well positioned to control network activity (Fig. 3a). We therefore used pre-embedding immuno-electron microscopy to examine Nav1.3 localization at MFTs in dorsal hippocampus of WT and 5xFAD mice (50), using a Nav1.3-selective antibody (51). Silver-intensified immunogold (SIG) labeling identified Nav1.3 in two distinct compartments of the mossy fiber-CA3 synapse; within MFTs (presynaptic) and within CA3 dendritic spines (postsynaptic) (Fig. 3b). Nav1.3 labeling in MFTs was significantly enriched in 3-month-old 5xFAD mice compared with age-matched controls and declined by 7 months (Fig. 3c). Thus, presynaptic Nav1.3 at MFTs is increased early in AD, following the same 3-month-high, 7-month-low trajectory as CA3 hyperactivity, supporting a role for presynaptic Nav1.3 expression at MFTs in driving hippocampal hyperexcitability in the prodromal 5xFAD mice.

**Figure 3.**
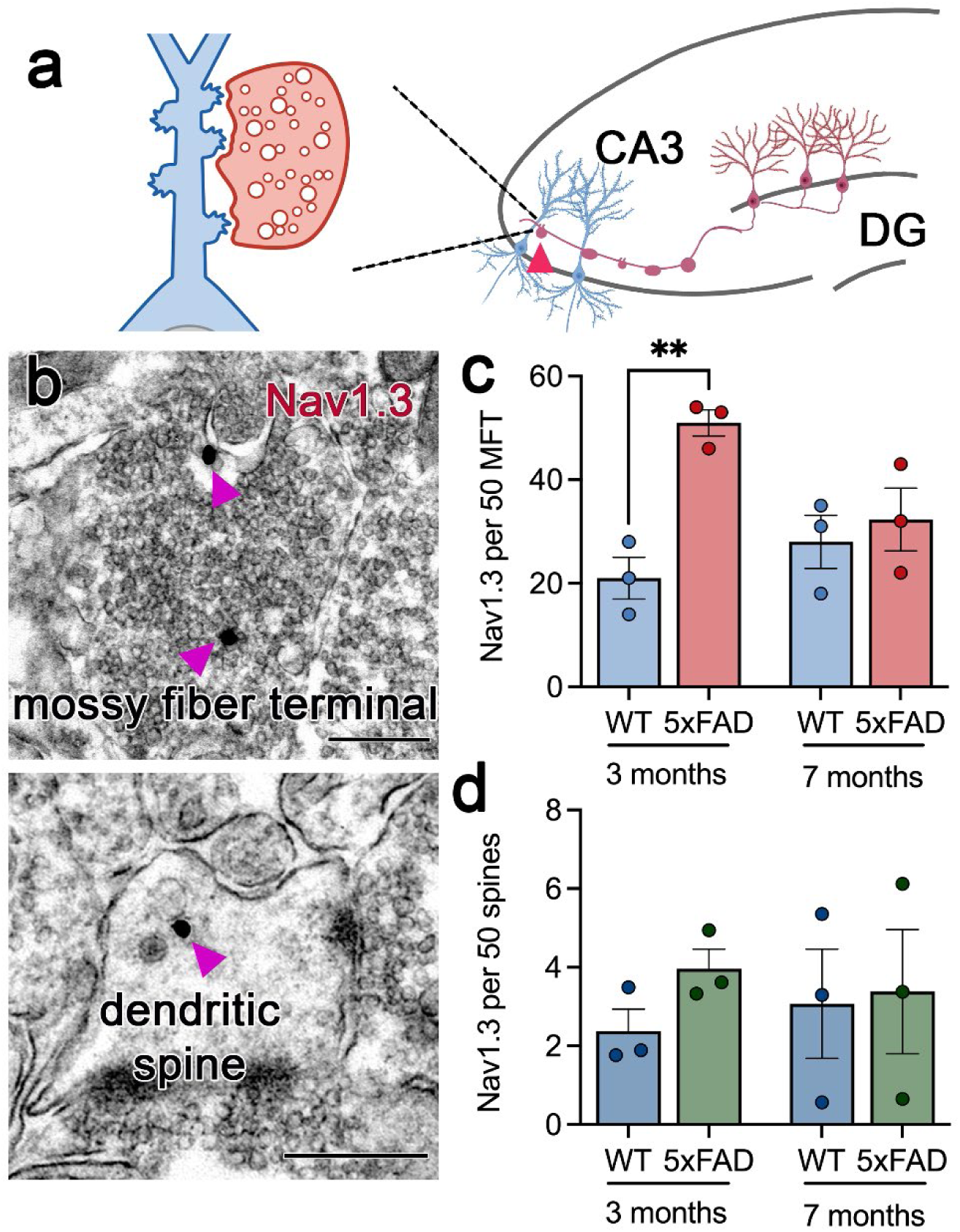
Nav1.3 expression is increased in mossy fiber terminals in early-stage 5xFAD AD-model mice. **a)** Schematic of mossy fiber terminal (red) contact (red arrowhead, right) on CA3 pyramidal cell dendritic spine (light blue). Inset (left) shows an enlarged representation of mossy terminal-CA3 spine contact **b)** Representative pre-embedding immuno-electron micrographs showing Nav1.3 in a mossy-fiber terminal (top, presynaptic) and a CA3 dendritic spine (bottom, postsynaptic). Purple arrowheads indicate silver-intensified immunogold (SIG) labeling for Nav1.3. Scale bar = 400 nm. **c)** Quantification of Nav1.3 SIG particles from 50 MFTs in WT and 5xFAD mice at 3 and 7 months. Data are mean ± SD; n = 3 mice per group. Two-way ANOVA revealed a significant genotype x age interaction (F(1, 8) = 7.68, p = 0.024). *Post-hoc* Sidak comparisons showed increased Nav1.3 SIG particles in MFTs from 3-month 5xFAD mice compared with 3-month WT mice (p = 0.007); no other pairwise comparisons reached significance. **d)** Quantification of postsynaptic Nav1.3 SIG particles per 50 CA3 spines per mouse per condition. Two-way ANOVA showed no significant effect of genotype p = 0.42, age p = 0.96, or interaction p = 0.58; n= 3 mice per group.

Nav1.3 was also present postsynaptically, in CA3 dendritic spines apposed to MFTs (Fig. 3b, d). Unlike the presynaptic compartment, spine Nav1.3 did not differ significantly by genotype or age (Fig. 3d). Nav1.3 immunogold was also observed on CA3 dendritic shafts, which were not included in the spine quantification. Together, these observations establish that CA3 pyramidal neurons contain a postsynaptic pool of Nav1.3, even though the disease-associated increase itself was most pronounced presynaptically.

### Knockdown of Nav1.3 reverses CA3 hyperexcitability in early-stage Alzheimer’s disease

Our data show Nav1.3 expression is increased within the early 5xFAD DG-CA3 circuit, in activated CA3 neurons (Fig. 2) and in mossy fiber terminals (Fig. 3). To test whether Nav1.3 contributes causally to hippocampal hyperexcitability in an AD model *in vivo*, we delivered a lentiviral vector encoding three *Scn3a*-targeting shRNAs (*Scn3a* encodes Nav1.3) into dorsal CA3 of 5xFAD mice and used a scrambled-shRNA virus as control. Efficient Nav1.3 knockdown in CA3 was confirmed by immunofluorescence (Fig. 4a-c); only mice with robust reduction in Nav1.3 expression and correct fiber placement were included in the analysis. For these experiments we focused on the 3-month time point and replotted the 3-month 5xFAD and WT data from Fig.1 alongside 5xFAD+scrambled and 5xFAD+Nav1.3-shRNA groups. Fiber photometry recordings showed that Nav1.3 knockdown markedly reduced CA3 neuronal activity in 5xFAD mice compared with both 5xFAD+scrambled and untreated 5xFAD mice (Fig. 4d). These results indicate that increased expression of Nav1.3 in CA3 contributes functionally to hyperexcitability, and that targeted reduction of Nav1.3 can attenuate abnormal network activity in the prodromal 5xFAD AD model.

**Figure 4.**
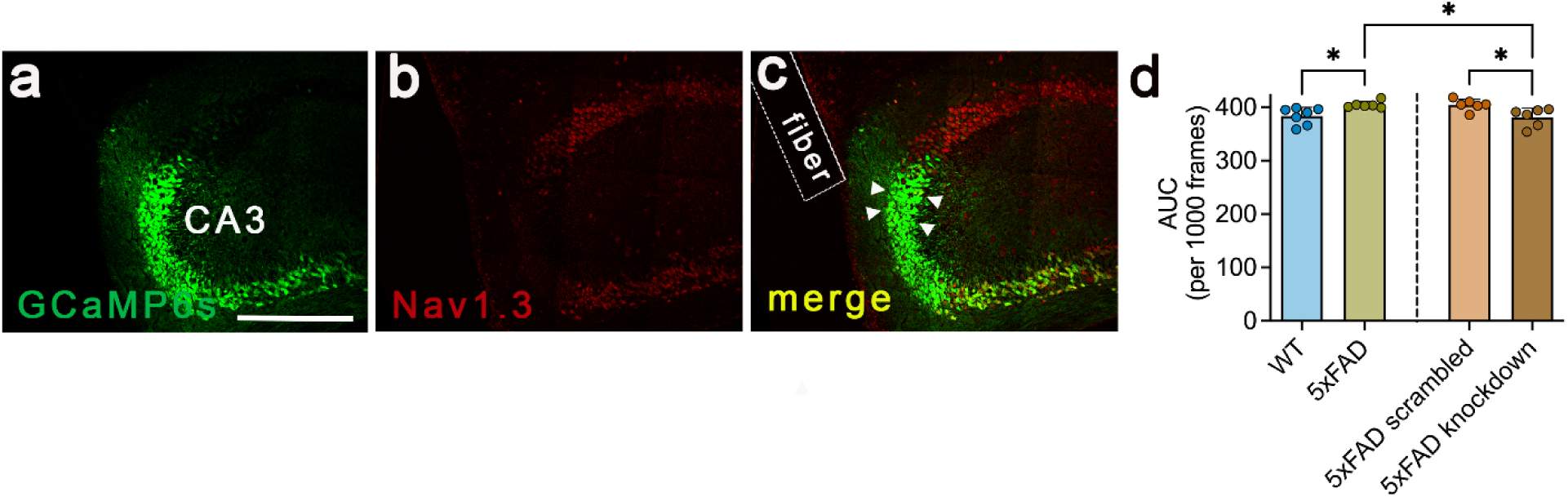
Nav1.3 knockdown in hippocampal CA3 rescues neuronal hyperactivity in early stage 5xFAD mice. **(a-c)** Representative confocal images showing GCaMP6s expression (green, a), Nav1.3 immunolabeling (red, b) and merged channel (yellow, c) in dorsal CA3 following viral delivery. Arrowheads indicate examples of GCaMP6s^+^ neurons with reduced Nav1.3 labeling after Nav1.3-shRNA; panel C also shows the implanted fiber tract (dashed outline) used for photometry. Scale bar = 400 μm **d)** Quantification of CA3 calcium activity (AUC/1000 frames) in 3-month-old WT, 5xFAD, 5xFAD+scrambled-shRNA, and 5xFAD+Nav1.3-shRNA mice. 3-month WT and 5xFAD data from Fig. 1c are replotted for comparison. Nav1.3 knockdown significantly reduced CA3 activity relative to both untreated 5xFAD and 5xFAD+scrambled controls. Data are mean ± SD; n = 6-7 mice per group. One-way ANOVA was used to assess treatment effects at 3 months (F (3, 21) = 5.63, p = 0.005), followed by post-hoc tests (Tukey) for pairwise comparisons: 5xFAD vs 5xFAD+Nav1.3-shRNA, p = 0.030; 5xFAD+scrambled vs 5xFAD+Nav1.3-shRNA, p = 0.032; WT vs 5xFAD, p = 0.047. WT vs 5xFAD+Nav1.3-shRNA, p = 0.987. WT and 5xFAD here are the Fig. 1c 3-month subset re-analyzed by one-way ANOVA, so the WT and 5xFAD p-value differs from the full two-way model in Fig. 1c).

While these results demonstrate that CA3-targeted Nav1.3 knockdown is sufficient to normalize CA3 hyperactivity, they do not resolve the synaptic compartment through which Nav1.3 acts. Nav1.3 expression is increased most prominently in presynaptic MFTs (Fig. 3c) yet is also present postsynaptically in CA3 spines (Fig. 3d), the compartment directly targeted by our CA3 knockdown. As the knockdown might not reduce presynaptic Nav1.3 within mossy fiber terminals, the rescue is most parsimoniously attributed to loss of postsynaptic CA3 Nav1.3, although a contribution from presynaptic Nav1.3 cannot be excluded. Definitive attribution will require compartment-specific manipulation or quantification of Nav1.3 at the postsynaptic mossy fiber-CA3 spine.

## Discussion

Developmentally expressed ion channels such as Nav1.3 are largely silenced in the mature brain, but whether their reactivation contributes to early AD circuit dysfunction has remained unexplored. Here, we used the 5xFAD mouse model to investigate the contribution of Nav1.3 to hippocampal CA3 hyperactivity. We show that increased CA3 neuronal activity coincided with increased expression of Nav1.3 early in disease progression, and that Nav1.3 knockdown attenuated this hyperactivity, establishing a causal role of Nav1.3 in mediating early hippocampal hyperexcitability. Notably, we observed these changes before the emergence of substantial Nab61^+^ Aβ oligomer accumulation, which is a pathological feature widely associated with AD progression (52).

Unlike Nav1.1 and Nav1.6, the principal voltage-gated sodium channels of the adult brain that have previously been implicated in AD-associated network dysfunction, Nav1.3 is predominantly expressed during early development and is largely absent in healthy mature brains (53, 54). In the adult nervous system, Nav1.3 is characteristically re-expressed under pathological conditions of hyperexcitability, following peripheral nerve and spinal cord injury, and in neuropathic pain and epilepsy (55, 56), yet its potential contribution to neurodegenerative diseases has received little attention. Our finding that Nav1.3 expression is increased in the early-stage 5xFAD hippocampus, where it contributes functionally to CA3 hyperexcitability, extends this injury-associated reactivation of a developmental excitability program to a model of AD.

Notably, elevated Nav1.3 expression preceded Nab61^+^ Aβ oligomer accumulation, indicating that increased Nav1.3 expression and the accompanying CA3 hyperactivity emerge before, and therefore cannot be a simple downstream consequence of, oligomeric Aβ. This temporal dissociation points to an oligomer-independent trigger acting early in disease. One candidate is beta-site amyloid precursor protein cleaving enzyme 1 (BACE1), whose activity is elevated in the AD brain (57). Beyond generating Aβ from APP, BACE1, together with γ-secretase, sequentially cleaves the auxiliary Nav β2 subunit, releasing a β2 intracellular domain that upregulates the mRNA and protein of pore-forming Nav α-subunits and thereby controls neuronal sodium current and excitability (57–59). This mechanism has been most directly established for Nav1.1 but raises the possibility that early rises in BACE1 activity similarly reactivate developmental Nav1.3, ahead of and independently of Aβ oligomer accumulation.

Nav1.3 expression at the DG-CA3 mossy fiber synapse is particularly consequential. The large MFTs of DG granule cells function as powerful ‘detonator’ synapses that can drive CA3 pyramidal cells to threshold with few or single presynaptic events, and they undergo pronounced short-term facilitation during repetitive activity (19, 20). Importantly, Navs are localized to both presynaptic boutons and postsynaptic dendritic spines (28), so Nav1.3 could shape excitability on either side of the synaptic cleft. A sodium channel with Nav1.3’s biophysical signature of low activation threshold, rapid recovery from inactivation, and support for sustained, high-frequency firing, is well suited to amplify this already potent drive at both compartments: presynaptically, by increasing the fidelity and gain of transmitter release, and postsynaptically, by boosting the local depolarization and spike output of CA3 neurons in response to that input. Because CA3 pyramidal neurons are further interconnected by an extensive recurrent collateral network, even a modest increase in excitatory drive can be recruited into network-level hypersynchrony (13). Increased Nav1.3 expression at this specific locus would therefore be expected to elevate CA3 population activity out of proportion to the same change elsewhere in the circuit, consistent with the early CA3 hyperactivity we observe. In this framework, the mossy fiber synapses are a site where Nav1.3 can exert leveraged control over hippocampal network state, on both sides of the synapse.

Beyond its immediate effect on network activity, early Nav1.3 increase might have longer-term consequences. Sustained, non-inactivating sodium influx imposes a persistent metabolic and ionic load: elevated intracellular Na^+^ can reverse the Na^+^/Ca^2+^ exchanger, promoting Ca^2+^ entry and dysregulation similar to that seen with neuronal injury in conditions of aberrant persistent sodium current (60, 61). Coupled with the broader evidence that chronic network hyperexcitability is not merely a correlate but an active driver of subsequent dysfunction and decline in AD (6), this raises the possibility that a transient early burst of Nav1.3-dependent hyperactivity acts as an initiating insult that leads to subsequent pathology. Such a mechanism could help explain the regional and cell-type specificity of neurodegeneration; populations that re-engage developmental excitability programs most strongly, or that are least able to buffer the resulting ionic and metabolic burden, could be rendered preferentially vulnerable, a pattern reminiscent of the circuit-specific neuronal loss that characterizes the human disease (62). The present findings establish Nav1.3 as a driver of early hyperexcitability and future studies will be necessary to directly demonstrate that this activity causes subsequent degeneration.

The functional normalization of CA3 activity following Na_v_1.3 knockdown raises the possibility that inhibiting Nav1.3 could represent a therapeutic strategy for hyperexcitability in early AD. Several Nav1.3 inhibitors have been developed, primarily for neuropathic pain, and recent structural studies have resolved antagonist-binding sites on human Nav1.3 (63, 64), providing a structural basis for subtype-selective drug design. Consistent with the druggability of this channel, we have shown that the pro-excitatory persistent and ramp currents of Nav1.3 are reduced by the volatile anesthetic sevoflurane (42), demonstrating that these hyperexcitability-associated currents are pharmacologically tractable. As Nav1.3 is largely absent from the healthy mature brain, a strategy targeting its pathological expression could in principle act selectively on the disease-associated channel pool while sparing sodium channels that mediate normal adult neurotransmission, potentially limiting the on-target central side effects that complicate broad Nav blockade. Their effects in neurodegenerative disease, however, have not been explored. Encouragingly, the recent approval of suzetrigine, the first selective Nav1.8 inhibitor, for acute pain demonstrates that subtype-selective sodium channel blockade is now clinically achievable (65). Notably, suzetrigine acts on a peripherally restricted channel, and CNS-directed subtype-selective Nav modulation remains to be established, having so far been limited by efficacy and off-target effects. Our findings nonetheless identify increased expression of Nav1.3 as a previously unrecognized target for reducing hippocampal hyperexcitability in early AD and motivate future work testing whether its pharmacological inhibition confers benefit in preclinical models and whether Nav1.3-mediated hyperexcitability contributes directly to cognitive decline and disease progression.

Several limitations of this study should be acknowledged. Our analysis focused on early-stage 5xFAD mice, and it remains to be determined whether increased Nav1.3 expression occurs in other amyloid or tau models, and, most importantly, whether it is recapitulated in human disease. We also characterized only a limited portion of the pathway; the upstream signals that reactivate Nav1.3 were not tested directly, and while knockdown normalized CA3 network activity, we did not measure whether this correction translates into improved cognition or altered disease progression. Determining whether Nav1.3-dependent hyperexcitability contributes causally to cognitive decline, and whether its suppression is beneficial, or given the physiological roles of network activity, potentially detrimental, will require longitudinal studies pairing Nav1.3 manipulation with behavioral and pathological endpoints. Finally, confirming that candidate Nav1.3 antagonists can engage the channel and dampen hyperexcitability *in vivo* would be an important step toward evaluating subtype-selective Nav modulation as a therapeutic strategy.

In conclusion, we identified increased expression of Nav1.3 as a previously unrecognized contributor to hippocampal CA3 hyperexcitability in early-stage 5xFAD AD model mice. We show that Nav1.3 is selectively upregulated in hyperactive CA3 neurons at the DG-CA3 mossy fiber synapse, and that reducing Nav1.3 expression restores CA3 network activity toward normal levels. These findings link increased expression of Nav1.3 to early AD-associated network dysfunction and support a broader view in which reactivation of immature excitability programs contributes to disease pathogenesis. More broadly, they raise the possibility that targeting Nav1.3 could limit pathological hyperexcitability during the early, potentially still-tractable window before extensive neurodegeneration occurs.

## Materials and Methods

### Animals

Hemizygous 5xFAD mice (B6.Cg-Tg(APPSwFlLon,PSEN1*M146L*L286V) 6799Vas/Mmjax, #034848-JAX) and wild-type C57BL/6J mice (JAX #000664) were obtained from The Jackson Laboratory (Bar Harbor, Maine) and bred in-house to maintain colonies. Animals were group-housed on a 12-hr light/dark cycle with ad libitum access to food and water. All procedures conformed to the National Institutes of Health Guide for the Care and Use of Laboratory Animals and were approved by the Weill Cornell Medicine Institutional Animal Care and Use Committee (IACUC). Animal handling and reporting adhered to ARRIVE guidelines.

### Viruses

AAV vector pAAV.Syn.GCaMP6s.WPRE.SV40 (Addgene #100843-AAV1) was purchased from Addgene (Watertown, MA) and the lentiviral vectors encoding Na^+^ channel Nav1.3 shRNA (sc-42647-V) or control shRNA (sc-108080) were obtained from Santa Cruz Biotechnology, Inc (Santa Cruz, CA).

### Materials

Ketamine hydrochloride was purchased from Covetrus (Portland, ME). Pentobarbital and other chemicals were obtained from Sigma-Aldrich (St. Louis, MO) or Covetrus. Isoflurane was purchased from Henry Schein Medical (Melville, NY). All remaining reagents and suppliers are specified as indicated.

### Antibodies

Primary antibodies used for immunofluorescence and immunogold-labeling were as follows. An anti-c-Fos antibody (rat polyclonal; SYSY #226017; Synaptic Systems, Göttingen, Germany) was used, raised against an N-terminal peptide sequence of c-Fos. An anti-Nav1.3 antibody (rabbit polyclonal; Alomone Labs #ASC-004; Jerusalem, Israel) was used, directed against an epitope in the C-terminal region of Nav1.3. The specificity of this antibody was characterized using the same criteria we previously applied to the Nav1.1, Nav1.2, and Nav1.6 antibodies (28). Nab61 is a mouse monoclonal antibody that selectively recognizes oligomeric Aβ species and was kindly provided by Dr. Virginia Lee (University of Pennsylvania); its characterization has been described previously (66). Secondary antibodies were species-specific goat anti-rabbit or goat anti-mouse IgG conjugated to Alexa Fluor dyes (Thermo Fisher Scientific, Waltham, MA): Alexa Fluor 488 (A-11034), Alexa Fluor 647 (A-21244, A21247), and Alexa Fluor 555 (A-21235).

### Stereotaxic surgery

5xFAD- and wild-type-mice (male and female, ∼P63, ∼P189) were anesthetized with ketamine (80 mg/kg) and xylazine (6 mg/kg), and placed in a stereotaxic frame (Kopf Instruments, Tujunga, CA). One drop of bupivacaine (2.5 mg/mL) was applied beneath the scalp for local analgesia prior to incision. Dexamethasone (2 mg/kg, i.p.) was administered intraoperatively to limit edema and meloxicam (2 mg/kg, i.p.) was administered intraoperatively for post-surgery analgesia. A craniotomy was made above dorsal CA3 (coordinates relative to bregma: AP -1.9 mm, ML +2.0 mm, DV -2.2 mm) and 200 nL of pAAV.Syn.GCaMP6s.WPRE.SV40 was injected unilaterally into CA3 at 50 nL/min using a Hamilton microliter syringe (7635–01) with a Nanofil needle (WPI NF33BV) that was left in place for 8 min after injection. For Na_v_1.3 knockdown, Na_v_1.3α shRNA lentiviral particles were mixed 1:1 with pAAV.Syn.GCaMP6s.WPRE.SV40 and injected at the same coordinates (total volume 500 nL). Scrambled-shRNA animals received scrambled shRNA lentiviral particles mixed 1:1 with pAAV.Syn.GCaMP6s.WPRE.SV40 (1:1 ratio; total volume 500 nL). Following viral delivery, a 400 μm-core optical fiber (2.5 mm length; Doric Lenses, Canada) was implanted above CA3 and secured to the skull with Vetbond Tissue Adhesive (3M, 1469SB) and Metabond dental cement (Parkell, S396). Mice were recovered on a warming pad, monitored until ambulatory, observed for 1 h post-op, and then returned to their home cages.

### *In vivo* fiber photometry

*In vivo* fiber recording was performed to measure calcium-dependent activity in freely moving animals as previously described (67). Three-to-four weeks after surgery, mice were habituated to a patch cord attached to the implanted optical fiber in an open field (40 × 40 cm) for 30 min; the patch cord length allowed unrestricted movement across the arena. Real-time Ca^2+^ transients were then recorded for 5 min using a fiber photometry system (Neurophotometrics MBF Bioscience, FP3001). Excitation at 415 nm (isobestic) and 470 nm was delivered through the implanted optical fiber implant via a 0.48 NA patch cord, and emitted fluorescence was returned through the same fiber, separated from excitation light by a dichroic mirror, and detected by an Orca-Flash4.0 V3 cMOS camera (Hamamatsu). Signals were digitized and acquired with Neurophotometrics software and preprocessed and normalized (ΔF/F) in Bonsai and MATLAB (R2026a, MathWorks) using custom scripts (see Supplementary File for code).

#### Calcium activity

Fiber photometry experiments were performed as previously described (68) in cohorts of n = 6-7 per sex. Animals with poor viral expression, incorrect fiber placement on histological verification, or clear outlier signals were excluded prior to analysis. Signals were acquired using Bonsai and analyzed offline in MATLAB. To correct for baseline drift, the isosbestic control channel was linearly fit to the calcium-dependent channel and subtracted. Relative fluorescence changes (ΔF/F) were then computed by defining a baseline fluorescence trace (F0) for each recording (average signal over a defined baseline window) and expressing each time point as the fractional change from this baseline ((F-F0)/F0). To facilitate comparisons across animals, ΔF/F values were converted to Z-scores by calculating the mean and standard deviation of ΔF/F during the baseline period for each animal and expressing each time point as the number of standard deviations above or below this mean. From these Z-scores traces, area under the curve (AUC), peak amplitude, and peak density within the analysis window were quantified using custom MATLAB scripts (Supplementary File).

### Nav1.3 immuno-electron microscopy

Electron microscopic immunolocalization of Nav1.3 was performed using the pre-embedding silver intensified immunogold method as described previously (50). Briefly, mice were deeply anesthetized with pentobarbital (150mg/kg; i.p.) and transcardially perfused with 5 mL 0.9% saline containing 2% heparin, followed by 30 mL of 3.75% acrolein and 2% paraformaldehyde (PFA) in 0.1 M phosphate buffer (PB; pH7.4). The brain was post-fixed in 2% acrolein and 2% PFA in PB for 30 min on a shaker. Tissue was then sectioned at 40 μm thickness with a vibratome (VT1000, Leica Microsystems) and stored in cryoprotectant solution at −20°C until further use. Three dorsal hippocampal sections per mouse were selected from each experimental condition and coded by hole-punches in the cortex. The sections were then placed in a single container and processed together through all immunocytochemical procedures to ensure identical labeling conditions. Sections were incubated in 1% sodium borohydride in PB for 30 min to neutralize reactive aldehydes, then rinsed in PB ∼10 times until gaseous bubbling ceased.

Sections were rinsed in 0.1 M Tris saline (TS, pH 7.6) followed by an incubation in 0.5% BSA diluted in TS for 30 min to reduce nonspecific labeling. sections were then incubated in rabbit anti-Nav1.3 antibody (1:50) in 0.1% BSA in TS at RT for 24 h and then 3 days at 4°C. Sections were rinsed in TS followed by washing buffer [0.02 M phosphate-buffered saline (PBS) with 0.8% BSA and 0.2% gelatin], and incubated overnight at 4°C in a 1:50 dilution of donkey anti-rabbit IgG conjugated to 1-nm gold particles [Electron Microscopy Sciences (EMS) Cat# 810.311] in 0.01% gelatin and 0.08% BSA. All primary and secondary antibody incubations were carried out at 145 rpm, and all rinses were conducted at 90 rpm on a rotator shaker. Sections were rinsed in PBS, post-fixed in 2% glutaraldehyde in PBS for 10 min, then rinsed in PBS and in 0.2 M sodium citrate buffer (pH 7.4). The conjugated gold particles were enhanced using silver solution (SEKL15 Sliver enhancement Kit, Prod No. 15718 Ted Pella Inc.) for 15 min. Sections were fixed in 2% osmium tetroxide for 1 h, dehydrated in increasing ethanol concentrations to propylene oxide, and embedded in EMBed 812 (EMS) between two sheets of Aclar plastic (Honeywell, Pottsville, PA) and left overnight. Following curing in an oven at 60–63 °C for 3 days, ultrathin sections (70 nm) of the CA3 were cut with a Diatome diamond knife (EMS) on an ultratome (Leica UCT6) and collected on 400 mesh thin-bar copper grids (T400-Cu, EMS). The grids were counterstained with Uranylless^TM^ (#22409) and lead citrate (#22410). Specimens were imaged on a Hitachi HT7800 transmission electron microscope.

#### Electron microscopy

Ultrathin sections from three mice per condition were examined at the tissue-plastic interface to minimize variability in antibody penetration (50). Immunolabeled profiles were classified using established morphological criteria (69). Mossy fiber terminals were identified by their characteristic ultrastructural features and large size (1-2 μm in diameter) and by their multisynaptic contacts onto dendritic spines of CA3 pyramidal dendrites (70). For quantification, 50 Nav1.3-labeled mossy terminals were sampled from randomly selected 55 μm^2^ fields within the CA3 region from each mouse and condition, and Nav1.3-immunogold particles within each terminal were counted manually

### Fluorescence immunohistochemistry

Mice were deeply anesthetized with sodium pentobarbital and transcardially perfused with heparin (100 mg/L) in PBS, followed by 4% PFA. Brains were rapidly dissected, hemispheres post-fixed in 4% PFA at 4 °C for 24 h, then cryoprotected in 30% sucrose in PBS at 4 °C. Coronal sections (40 μm) were cut on a vibratome (VT1000, Leica Microsystems) and stored in cryoprotectant (30% sucrose, 30% ethylene glycol in PB) at -20°C until use. For immunofluorescence, free-floating sections were rinsed with PBS and permeabilized in PBST (PBS containing 0.2% Triton X-100). Sections were then blocked in PBST containing 2% bovine serum albumin (BSA) for 60 min at RT with gentle agitation, followed by overnight incubation at 4 ℃ with rabbit anti-c-Fos (1:500), rabbit anti-Nav1.3 (1:200), or mouse anti-Nab61 (1:500) diluted in PBST/2% BSA. After rinsing in PBST, sections were incubated with secondary antibodies (1:500) for 60 min at RT, rinsed again in PBST, and mounted on coated slides with ProLong Diamond Antifade Mountant (Thermofisher, P36961). No antigen retrieval was performed.

### Confocal microscopy

Immunofluorescence labeling were captured using a 20x/0.8 NA Plan-Apochromat air objective (Zeiss, Thornwood, NY) on a Zeiss Axio Observer Z1 microscope equipped with a CSU-X spinning disk confocal scanner (Yokogawa, Wayne, PA). Samples were excited with a laser launch equipped with 50 mW solid-state lasers (488 nm, 561 nm), and fluorescence emission was collected through 525/50 or 629/62 band-pass filters. Images were acquired using an Orca Flash4 camera (Hamamatsu, Bridgewater, NJ) and Zeiss Zen Blue software.

#### Fluorescence microscopy

Aβ oligomer expression was quantified using Nab61 immunolabeling (71). Integrated density of Nab61 was measured in three randomly selected, non-overlapping fields (200 x 200 μm) within the CA3 region per animal, averaged, and then normalized to area to yield density per 100 μm^2^. Nav1.3 and c-Fos expression were quantified by counting c-Fos^+^ cells per 1000 μm^2^ and scoring Nav1.3-positive puncta colocalized with the soma of those c-Fos-positive cells. Images were analyzed using Fiji/ImageJ (NIH, version 1.54p), and CA3 boundaries were defined according to Allen Brain Atlas. Sample sizes for immunohistochemistry experiments were estimated based on previous literature (72), with cohorts of n=3 per group.

### Statistical analysis

All analyses were performed by investigators blinded to the experimental group. Statistical comparisons were conducted using unpaired two-tailed Student’s t test or one-way/two-way ANOVA, followed by Šídák’s or Tukey’s post-hoc tests, as appropriate, using GraphPad Prism (version 11.0.1; San Diego, CA, USA). A significance threshold of p < 0.05 was applied. Data are presented as mean ± SD. Asterisks denote values significantly different from control: \**P* < 0.05, \*\**P* < 0.01, \*\*\**P* < 0.001, \*\*\*\**P* <0.0001. Sample size calculations were refined using interim power analysis (α = 0.05, power = 0.8) based on preliminary effect sizes, and additional animals were added accordingly. Sample sizes are given in the figure legends; n refers to individual mice. CA3 photometry data were analyzed by full-model two-way ANOVA with genotype (Wt, 5xFAD) and age (3, 7 months) as between subjects factors, performed on the pooled cohort and separately within each sex. Two comparisons were pre-specified as planned contrasts based on our a priori hypotheses, 3-month WT vs 3-month 5xFAD, and 3-month vs 7-month 5xFAD, and were evaluated with Šídák’s correction for the two planned comparisons. The same analysis was applied identically to the pooled and sex-stratified data.

## Acknowledgments

We thank Dr. Virginia Lee (University of Pennsylvania) for generously providing the Nab61 antibody. We also thank Ms. June Chan of the Neuroanatomy EM core, Feil Family Brain and Mind Research Institute, for technical assistance.

## Author contributions

JP and JT conceived and designed the study. Material preparation, data collection, and analysis for Figures 1-4 were performed by JT. KP and JX contributed to data collection for Fig. 3. AJ contributed to data collection for Fig. 4. KP contributed to data analysis for Fig. 3. CS contributed to data collection for Figs. 2 and 4 and assisted with figure formatting across all figures. Custom macros were written by CS. Figures were prepared by JT, JP, and CS. The original draft of the manuscript was written by JT and JP, with additional review and editing by JP, HCH, and TAM. All authors approved the final manuscript.

## Supporting Materials

**Supplementary Figure 1.**
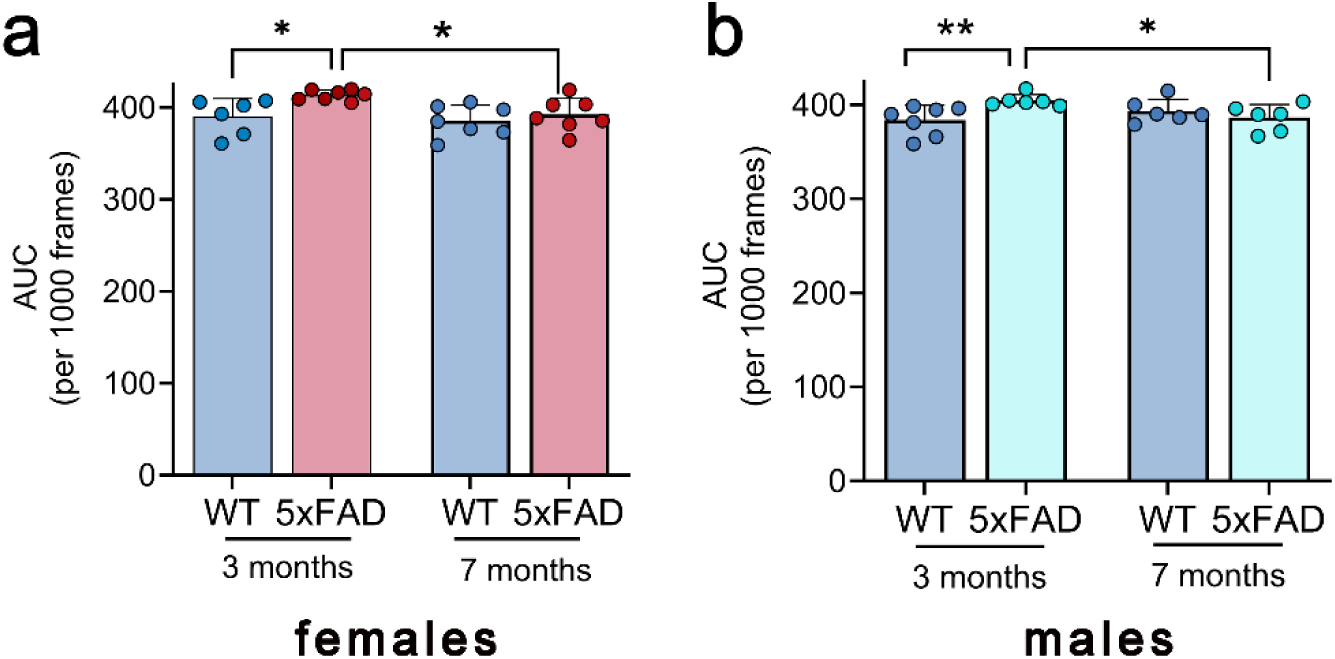
CA3 hyperactivity is present in both sexes. **a-b)** Quantification of CA3 GCaMP6s photometry (Area under the curve (AUC)) for females (a; n = 27) and males (b; n = 25) separately. Data are mean ± SD. Full-model two-way ANOVA (genotype x age) was performed separately for each sex, with two pre-specified planned comparisons (3-month WT vs 3-month 5xFAD; 3-month vs 7-month 5xFAD) evaluated using Sidak’s correction. In females (a), there were significant main effects of genotype (F(1, 23) = 6.08, p = 0.022) and age (F(1, 23) = 4.47, p = 0.046) without a significant genotype x age interaction (F(1, 23) = 1.86, p = 0.186). In males (b), there was a significant genotype x age interaction (F(1, 21) = 7.23, p = 0.014) without significant main effects of genotype (F(1, 21) = 1.85, p = 0.189) or age (F(1, 21) = 0.68, p = 0.419). In both sexes, planned comparisons confirmed higher CA3 activity in 3-month 5xFAD than 3-month WT (females, p = 0.014; males, p = 0.008) and than 7-month 5xFAD (females, p = 0.020; males, p = 0.024). Both sexes thus show the same directional pattern of early CA3 hyperactivity, expressed through significant genotype and age main effects in females and a significant genotype x age interaction in males.

### Supplementary Method

#### MATLAB script for fiber photometry signal processing and quantification of Z score

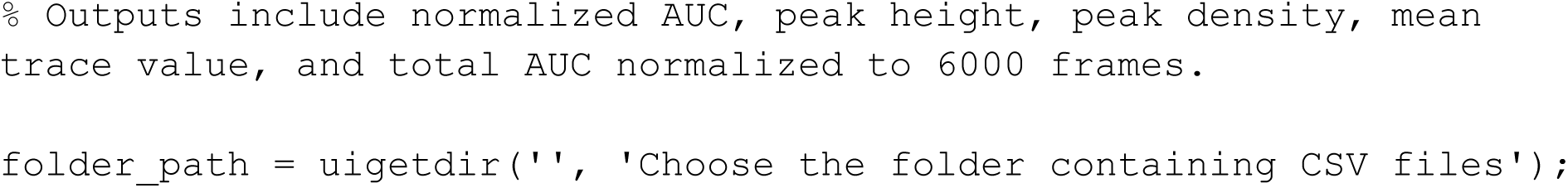
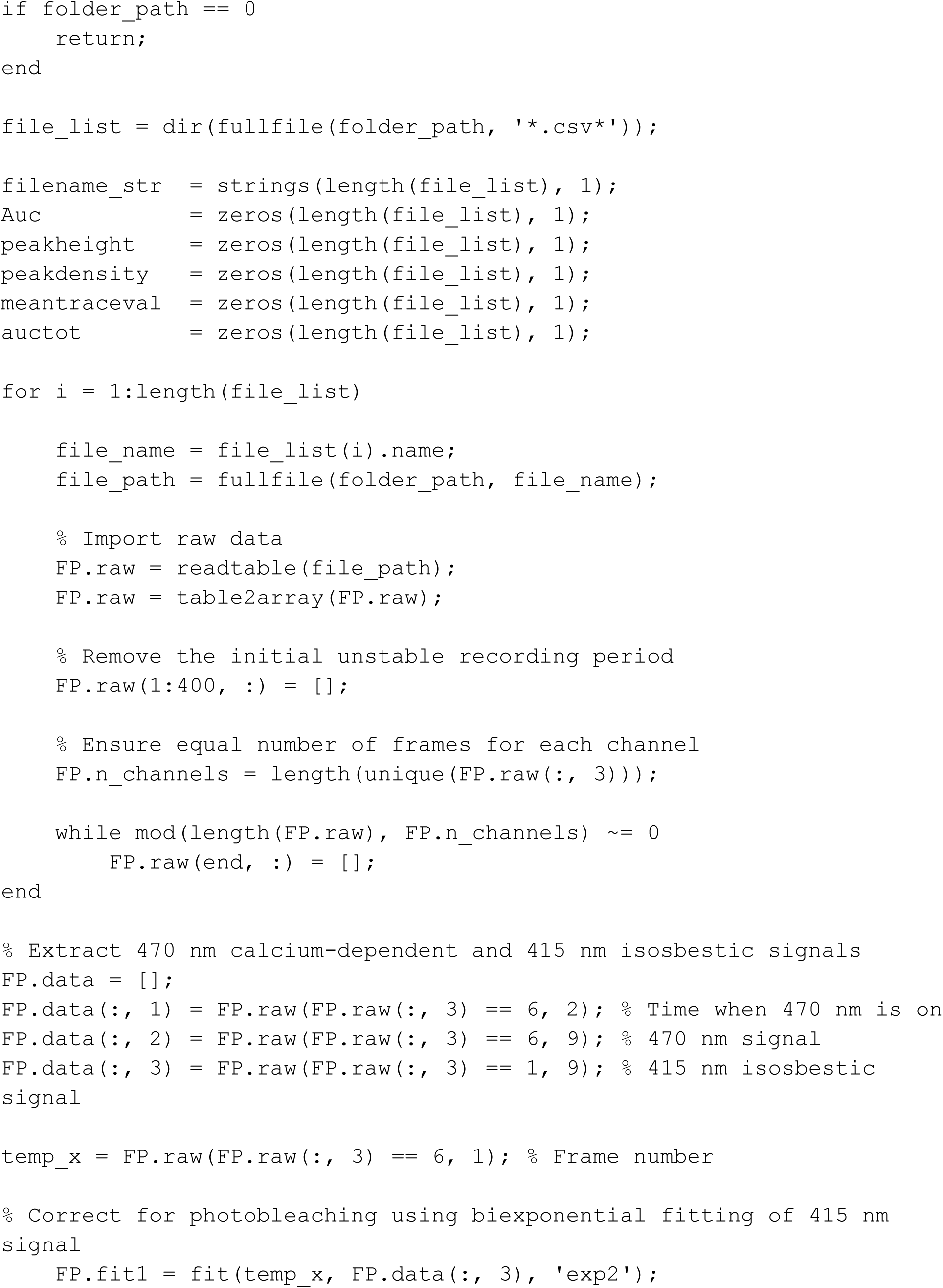
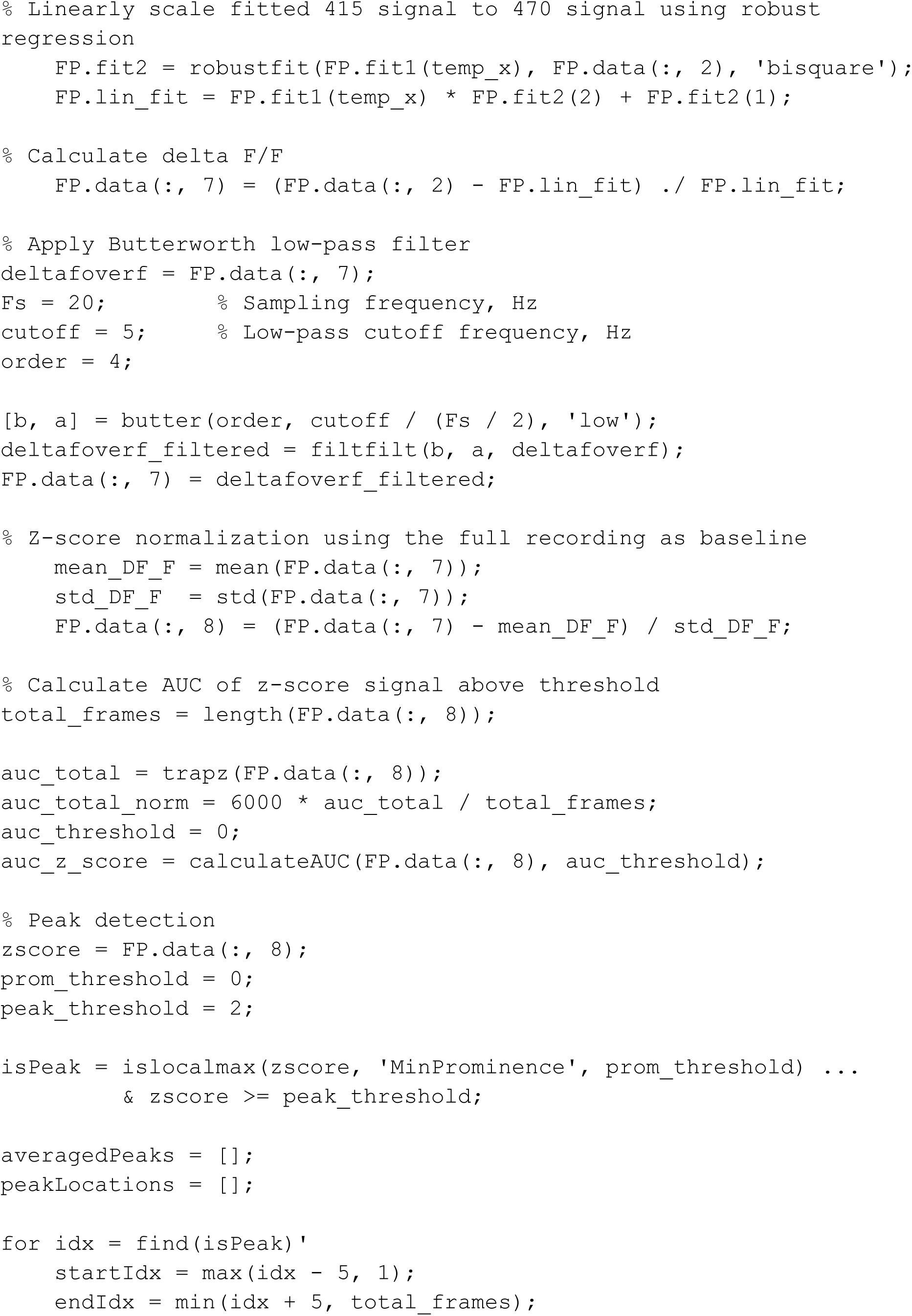
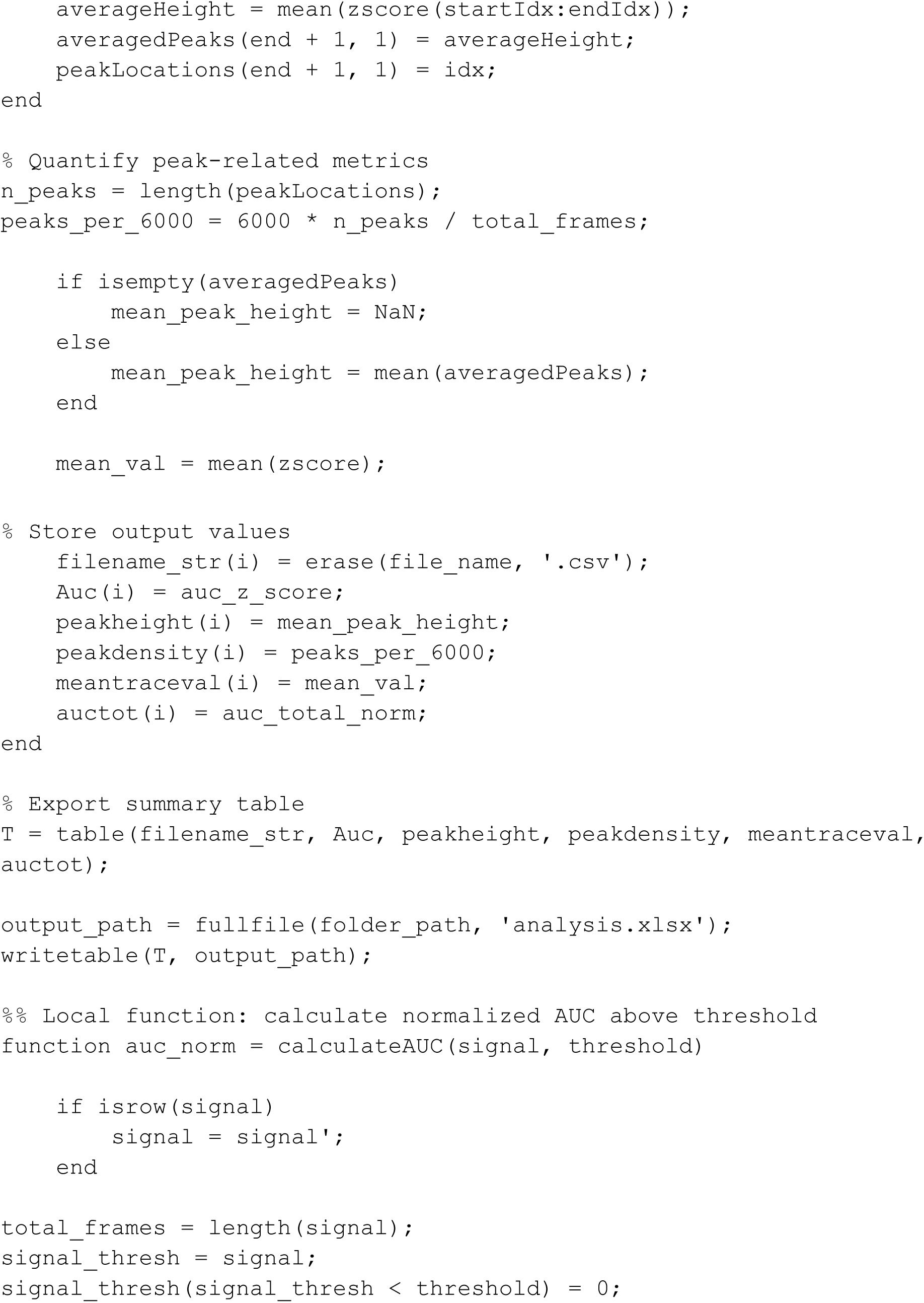
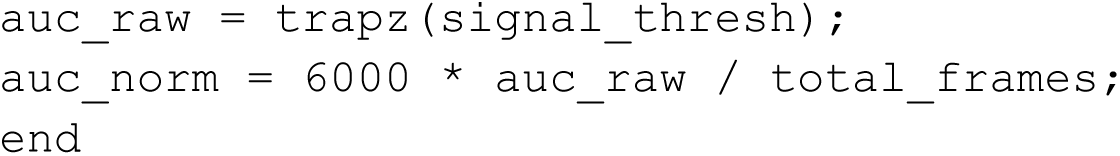

